# evopython: a Python package for feature-focused, comparative genomic data exploration

**DOI:** 10.1101/2023.09.02.556042

**Authors:** Steven T. Mick, Ana Fiszbein

## Abstract

**Motivation:** Software such as *LiftOver* allows for the intra- and inter-species conversion of genomic coordinates between genome assemblies, but this coordinate-centric workflow is naive to sequence alterations underling the conversion. For instance, it does not clarify the location of any insertions or deletions that might have occurred.

**Results:** To facilitate inter-species, sequence-based analyses, we developed *evopython*, a simple, object-oriented Python package that enables the sequence-aware resolution of genomic coordinates directly from pairwise and multiple whole-genome alignment data. The output is a Python dictionary storing all information relevant to the alignment: the participating species’ names, the position of the alignment in each species’ genome assembly, and the aligned sequences.

**Availability and implementation:** The source code and documentation are available at https://github.com/fiszbein-lab/evopython.

## Introduction

Darwin’s profound insight, universal common descent, involved reframing the apparent similarities between species as evidence of shared ancestry. This insight has developed into a fundamental concept that we employ to compare genomes and annotate genomic features, stretches of DNA that may have a specific function.

The alignment of sequences sampled from different species serves as initial evidence of identity by descent, an indication that these species may have descended from a common ancestor (Gillespie 1998). At a larger scale, alignment data can be utilized to generate estimates of conservation across the genome, highlighting sequences that are under purifying selection and likely to be functionally important (Cooper *et al*., 2005). This comparative framework has been leveraged to annotate genomes (Alexandersson *et al*., 2003; King *et al*., 2005). Furthermore, because sequence conservation often implies functional conservation, it is an invaluable resource for molecular biology, enabling the exchange of hypotheses regarding molecular phenotypes across different species.

It’s become increasingly tractable to sequence and assemble entire genomes, and popular databases (Herro *et al*., 2016; Nassar *et al*., 2022) have kept pace to deliver pairwise and multiple whole-genome alignment data for broad ranges of species. Moreover, functional genomic techniques continue to generate complementary molecular phenotype data, allowing direct annotation—rather than sequence-based inference—of genes and other relevant features that comprise functional genomes. Although there is much insight to be gained where whole-genome alignment and molecular phenotype data overlap, it can be cumbersome to integrate the two in a manner that is conducive to holistic exploration.

To support such analyses, we developed *evopython*, a free and open-source Python package that reconciles genomic features with whole-genome alignment data, providing an interface for investigating the former within the context of the latter. While popular software such as LiftOver (Kuhn *et al*., 2012) enables the intra- and inter-species conversion of coordinates between genome assemblies, it lacks insight into sequence alterations during conversion. On the other hand, *evopython* allows for direct, feature-focused access of whole-genome alignment data, providing both the position and sequence of each alignment. Here, we implemented *evopython* to study the sequence divergence in a human core promoter and the link between splice-site mutation and isoform count changes across species. *evopython* accommodates the nested and contingent boundaries inherent to genomic features with dictionary-like data structures that combine feature-wise iteration and key access with sense-aware coordinate manipulation for flexible generation of derived features.

## Description

In genome-scale sequence analyses, it’s standard to represent features of interest, such as splice sites or transcription factor binding sites, with their genomic coordinates. The actual sequences are then retrieved only after any required coordinate manipulation or conversion has been executed. A substantial amount of popular software is dedicated to these coordinate-centric analyses (e.g., Dale *et al*. 2011 and Lawrence *et al*., 2013), but sequence-naive approaches do not translate often well to a comparative genomic context.

*LiftOver* (Nassar *et al*., 2022), for example, is a popular tool that can be used to interface with pairwise, whole-genome alignment data: coordinates from one species’ reference genome can be converted to coordinates in another species’ reference genome. *BedTools* (Quinlan and Hall, 2010) can then be used to derive sequences, but because retrieval is from ungapped references, the actual alignment is ambiguous. *evopython*, on the other hand, provides direct access to both pairwise and multiple whole-genome alignment data, so while querying is still coordinate-based, both the positional and sequence information of the alignment are retrieved (Fig. 1).

**Fig 1.**
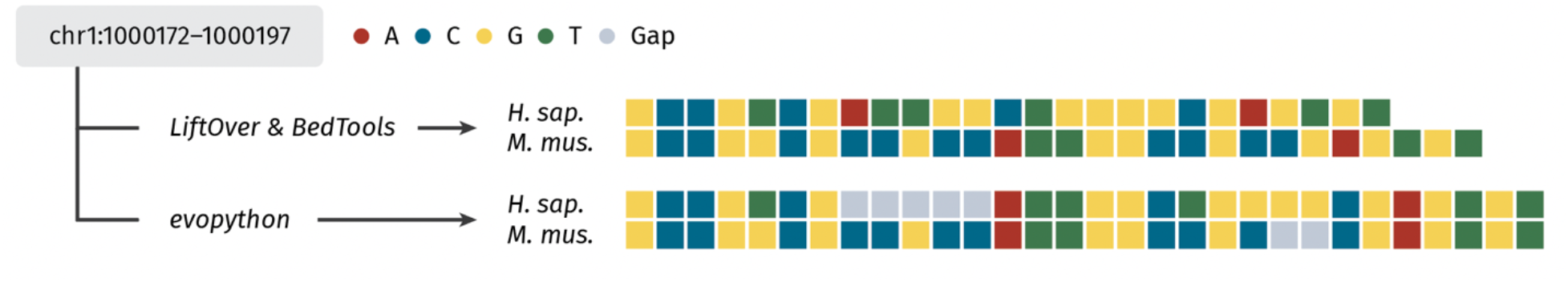
A comparative genomic workflow with *LiftOver-BedTools* and *evopython*. Given some human genomic interval, we can use *LiftOver* to convert the coordinates to the mouse genome assembly, effectively retrieving the positional alignment, and *BedTools getfasta* to then obtain the sequences from either species’ reference. This *LiftOver-BedTools* approach, however, does not report how the sequences align—where and in which assembly any gaps are located. The *evopython* workflow, on the other hand, provides a direct interface to the whole-genome alignment, thus allowing retrieval of the alignment’s structure in addition to the genomic position. *H. sap, Homo sapiens; M. mus*., *Mus musculus*

*evopython* is a modular, object-oriented Python package, specifically designed for parsing features at genome-scale and resolving their alignments from whole-genome alignment data (Fig. 2). The core parser classes inherit from the *UserDict* class, a container specified in the *collections* standard library. *UserDict* enables child classes to simulate native dictionaries and achieve similar behavior. To provide finer control over the representation of input features, we implemented a custom _*init*_ method for each parser. As a result, the generated objects are custom data structures that retain the familiarity of native dictionaries and can be interacted with accordingly. The parsing process aims to converge on instances of the *Feature* class, which serves as a data container for representing stranded genomic intervals. Instances of the *Feature* class are hashable and support sense-aware coordinate manipulation, including options for 5’ or 3’ centering. These functionalities facilitate the collection of unique and contingent features, enabling comprehensive analysis within the context of genomic structures.

**Fig 2.**
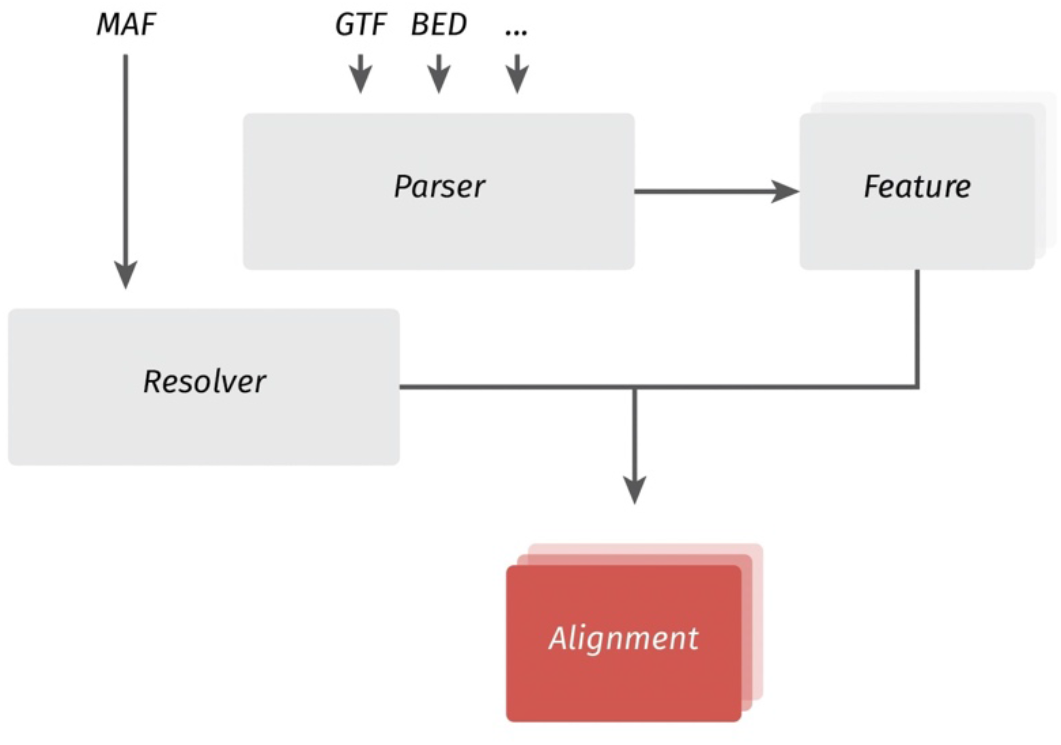
*evopython*’s modular, object-oriented design. The fundamental capabilities of *evopython* are encapsulated within two key class-level functionalities: Parser and Resolver. The Parser class provides a dictionary-like interface for interacting with feature-storing formats, such as GTF or BED. Through this interface, accesses are directed towards *Feature* instances, which represent the parsed features. The Resolver class then resolves these features from within the context of the whole-genome alignment. It performs the task of mapping the features onto the alignment and returns a nested dictionary representation that reflects the alignment structure.

**Fig. 3.**
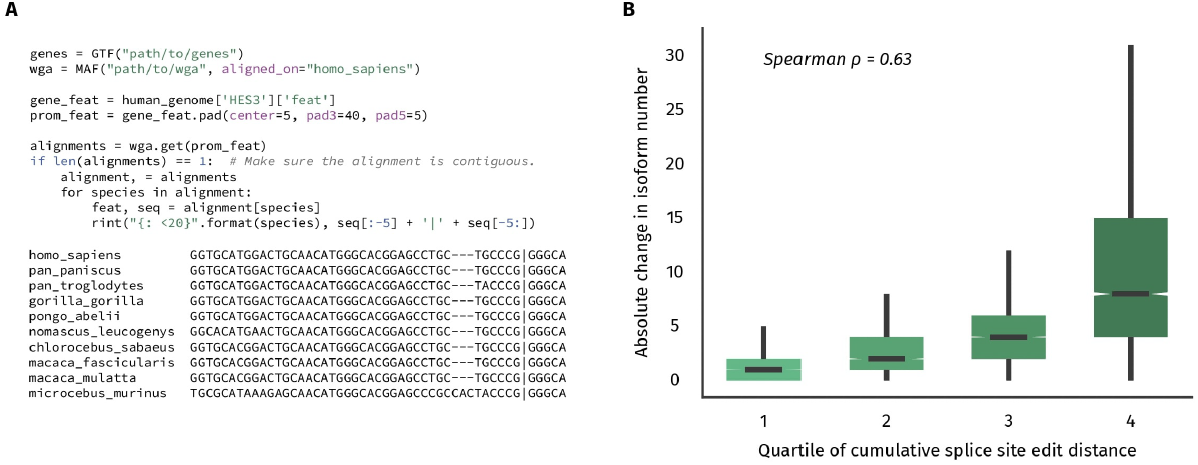
Usage of evopython for gene- and genome-scale analyses. **(A)** Genes can be accessed via name and secondary features derived using the *pad* method. The derived feature can then be resolved from within the whole-genome alignment and formatted for visualization, as we do here for the human transcription factor HES3 and 10 primates multiple whole-genome alignment dataset. **(B)** It’s also possible to derive secondary features at genome-scale and further incorporate gene annotation information. Here, we focus on human protein-coding genes with protein-coding orthologs in the mouse genome, exploring the relationship between the length-normalized edit distance between aligned human and mouse splice sites and the absolute difference in their host genes’ numbers of annotated isoforms.

Alignment querying is handled with the *MAF* class, with each *Feature* instance being passed to the *get* method for resolution. *get* handles the wide range of cases that can impede resolution and returns a nested dictionary representing the alignment. It is worth noting that *evopython* supports partial resolution. In cases where a feature is not entirely contained within a single alignment record or when a feature is discontinuous, spanning multiple records, *evopython* is equipped to handle such scenarios. It dynamically adjusts the feature by contracting or breaking it up as necessary, resulting in alignment records that accurately represent the modified feature structure.

To ensure accuracy, we generated 100 random features from 10 pairwise and multiple whole genome alignment datasets, testing that the resolved alignment—the sequences and coordinates for each participating species—matched the corresponding Ensembl alignment, retrieved from their REST API (Yates *et al*., 2014). Testing revealed no discrepancies (see Supplementary Table 1 for meta-data and the GitHub page for code).

The core *evopython* package was developed with Python 3.10.8 and relies on just the well-supported *biopython* package (Cock *et al*., 2009). The testing module additionally uses *requests* (Reitz 2023) to interact with the Ensembl REST API.

## Usage and documentation

*evopython* can be installed from the command line with *pip install evopython*. To demonstrate its usage, we use a multiple whole genome alignment dataset comprising 10 primates and the pairwise human-mouse whole genome alignment, both downloaded from Ensembl (Herro *et al*., 2016). Complete documentation and Jupyter notebooks to reproduce the forthcoming analyses are available on our GitHub page: https://github.com/fiszbein-lab/evopython.

Suppose we wanted to explore the sequence divergence in the human HES3 transcription factor’s core promoter, given the orthologous gene in most primates has a transcription start site further downstream. Here, we can leverage the fact that some eukaryotic gene features are defined from their transcription start site to derive the region upstream. The resultant core promoter-associated feature is resolved from within the whole-genome alignment and returned as a dictionary, from which we can produce a neatly-formatted representation in standard output (Fig. 2A).

The power of *evopython* is best demonstrated when integrating both sequence and molecular phenotype information: if more than one species comprising a pairwise or multiple whole genome alignment has well-annotated genes, for example, we can incorporate that information into our analyses. Suppose we wanted to explore the connection between splice-site mutation and changes in isoform number since the human and mouse lineages diverged. Using extant orthology data (The Alliance of Genome Resources Consortium 2019), we can iterate over the human gene set; check whether each gene is both protein-coding and has a protein-coding ortholog in the mouse genome; and derive the set of unique splice sites from the genes’ constituent exon features in a manner analogous to the above core promoter derivation. Then, we can resolve these sites from the whole-genome alignment; sum the edit distances between the aligned human and mouse sequences, normalizing on the splice site length; and visualize the relationship between cumulative splice site edit distance and the host gene’s absolute change in isoform number—an attribute obtainable from both species’ annotation sets (Fig. 2B).

## Discussion

While there’s a considerable on-going effort to develop packages for phylogenetic analyses such as tree inference (e.g., Dylus *et al*., 2023), the basic task of feature alignment retrieval at genome-scale has remained a challenge. *evopython*, to that end, provides a flexible and intuitive interface for feature-focused comparative genomic data exploration. Because the package is modular in structure, with feature collection separated from alignment querying, it can be readily extended to support the range of formats used to represent genomic features. We anticipate *evopython* will make genome-scale, feature-centric comparative genomic analyses more accessible for the growing collection of species with both sequenced and functionally-profiled genomes.

